# Integrating complexity in population modelling: from matrix to dynamic models using the jellyfish *Carybdea marsupialis*

**DOI:** 10.1101/2024.01.15.575756

**Authors:** Adrián Flores-García, John Yañez-Dobson, Eva S. Fonfría, David García-García, César Bordehore

## Abstract

Matrix models are widely used in population ecology studies and are valuable for analysing population dynamics. Nonetheless, this approach is somewhat rigid in terms of generating complex scenarios. Starting from the values of the transition matrix, we can build a dynamic model to incorporate more biological-based reality into the model (e.g. polyp stage) and provide a higher flexibility in generating scenarios. As an example, we used the transition matrix calculated for a time series data of a population of the box jellyfish *Carybdea marsupialis* (L. 1758) in the Western Mediterranean in a previously published study. Dynamic models can help us to better understand the complex relationships that drive populations, test different hypotheses and compare scenarios. The dynamic model was developed in STELLA Architect, calibrated and optimised and it has been used to simulate various scenarios of ecological interest, including a decline in food supply, jellyfish removal strategies, changes in drift currents and changes in substrate availability for planulae to settle. A sensitivity analysis showed that polyp strobilation rate and strobilation pattern were two of the most sensitive variables. This matrix-to-dynamic model approach could be useful to integrate more biological complexity into population models and, in turn, obtain a better fit to the field data.

## INTRODUCTION

The study of population dynamics typically focuses on the variation of individuals’ abundances within each age or size class over time and space. Descriptive studies of the evolution of a population have limited usefulness since a descriptive approach cannot disentangle non-linear relationships, e.g. growth and death rates *vs* abundances at each size class. A matrix approach can be used to construct useful models to get insight into the dynamics, thus based on age classes (Leslie matrix models) or stage/size classes (Lefkovitch models). These models have been widely used in population dynamics studies and they are useful tools to analyse the stability of the population, the stable structure, the reproductive value of each class and to perform different analyses, such as sensitivity analysis, or test different hypotheses for both retrospective and prospective studies (Caswell, 2001; Crone et al., 2011). However, matrix population models have some constraints, such as a low flexibility in the design that precludes optimal adaptation to the complexity of biological cycles, and limitations of matrix algebra.

Bordehore et al. (2015) calculated the parameters of the transition matrix for a population of *C. marsupialis* (Linnaeus 1758) (Cubozoa) using a time-series data of density and size. Since most swimming marine organisms cannot be technically tagged and followed, in order to obtain survival rates over time, the matrix parameters were obtained through an approach known as “inverse problem” (Wood, 1997). This method has only been used rarely, e.g. for *Octopus vulgaris* (Katsanevakis & Verriopoulos, 2006) and for an invasive aquatic plant *Alternanthera philoxeroides* (Erwin et al., 2012).

Dynamic population models have been used in population modelling for some marine species, such as amphipods (Anastácio et al., 2003; Gamito et al., 2010; Martins et al., 2009), barnacles (da Silva et al., 2009), bivalves (Hawkins et al., 2002), brachiopods (Valentin & Marazzo, 2003), northern fur seals (Lee et al., 2014), fishes (Mahévas & Pelletier, 2004; Yamashita et al., 2016; Bala et al., 2014), copepods (Marín, 1997), Kemp’s ridley sea turtle (Kocmoud et al., 2019), aquatic macroinvertebrate communities (Gertseva et al., 2004) and aquatic plants (Jarvis et al., 2004). To the best of our knowledge, no known dynamic model has been developed for any cubozoan species.

The Class Cubozoa (Phylum Cnidaria), also known as box jellyfish, comprise around 50 species (Collins & Jarms, 2022) in two orders, Chirodropidae and Carybdeidae. They are found in coastal waters worldwide mainly in tropical and temperate waters (Kingsford & Mooney, 2014). *C. marsupialis* is one of the 21 species in the Carybdeideae family (Collins & Jarms, 2022) and the most abundant of the only two species described for the Mediterranean Sea (Acevedo et al., 2019); the other species is *Copula lucentia sp.nov.* (Fonfría et al., *in press*).

Cubozoans are strong active swimmers (Garm et al., 2007), but their swimming ability is limited during small juvenile phases. As they grow larger, their swimming ability enables them to overcome weak currents (Bordehore et al., 2023), adjust their position in the water column, and potentially select habitats that have a higher food availability (Bordehore et al., 2014). The food preferences of cubozoans vary with their size; smaller individuals prey on small planktonic organisms such as rotifers, copepods or brachiopods, while larger mature individuals feed on brachiopods, other larger crustaceans and even fish (Acevedo et al., 2013). There is no data available about known predators of *C.marsupialis*. Even *Caretta caretta* (Revelles et al., 2007), one of the Mediterranean sea turtles known to prey on jellyfish, has not been shown to be an active predator of *C. marsupialis* (C. Bordehore pers. comm.). Only predators of *Chironex fleckerii* have been described, such as the sea turtle *Chelonia mydas* (Hamner et al., 1995) and fish species such as *Scomberomorus commerson, Nemadactylus valenciennesi* and *Oplegnathus woodwardi* (Barnes & Kinsey, 1986).While the abundance of cubozoans in coastal waters is generally low and uneven (Kingsford & Mooney, 2014), there are certain locations where their density can reach high numbers. Examples include *Alatina moseri* in Hawaii, *Alatina alata* in the Caribbean Netherlands and *Alatina mordens* in Australia (Lawley et al., 2016), as well as *C. marsupialis* on the south-western Spanish coast, northeast Atlantic (Rodríguez-García et al., 2021) and the Western Mediterranean, where they can be responsible for most of the stings that occur on some beaches where this species is present as a resident species since it is not advected by currents in its adult stage (Bordehore et al., 2011; Bordehore et al., 2020).

The sting of cubozoans can range from mild to severe symptoms for humans, depending on the species and the patient’s reaction to the venom (Fenner et al., 2010; Lippmann et al., 2011). Some species, such as *C. marsupialis* (Peca et al., 1997) or *Alatina alata* (Crow et al., 2021), cause only mild stings leading to local pain, itching, and redness on the epidermis, while others, like *Chironex fleckeri, Carukia barnesi* (Fenner, 2005), *Chiropsoides buitendijki* (Lippmann et al., 2011), and *Morbakka fenneri* (Gershwin, 2008), can be lethal under certain circumstances. The sting of *C. marsupialis* can also lead to systemic effects on sensitive individuals (Bordehore et al., 2015).

This work transforms a matrix model of *C. marsupialis*, previously published in Bordehore et al. 2015, into a dynamic model, by using calculated projection matrix parameters, adding more biological complexity, such as the benthic phase (polyps) and optimising the strobilation pattern. This approach intends to obtain a better fit of the output of the model with the observed data. It also allows us to perform different scenario comparisons and sensibility analysis on some of the parameters that rule the model.

## METHODS

### General characteristics of the model

We used the STELLA Architect 2.0.1 (www.iseesystems.com) program. Dynamic models are created with the combination of stocks, flows, converters, and connectors. Stocks accumulate individuals, flows are used to fill or drain stocks, and converters are used to set rate values (which can be constant or variable). The mathematical relationships between the different parts of the model are described inside flows or converters.

We used the data from Bordehore et al. (2015), where abundances for each size class were obtained using net sampling along the coast of Denia (38° 51’ 59.9’’N, 0° 08’36.6’’O), south-east Spain, from June 2010 to October 2011, comprising an entire planktonic cycle. The medusa phase was split into 6 planktonic stages (Juveniles 1 to 3 and Adults 1 to 3 – see Table 1 for biological criteria and Table 2 for the abundance of each size class) and growth rates were obtained from the following transition matrix, calculated by Bordehore et al. (2015):

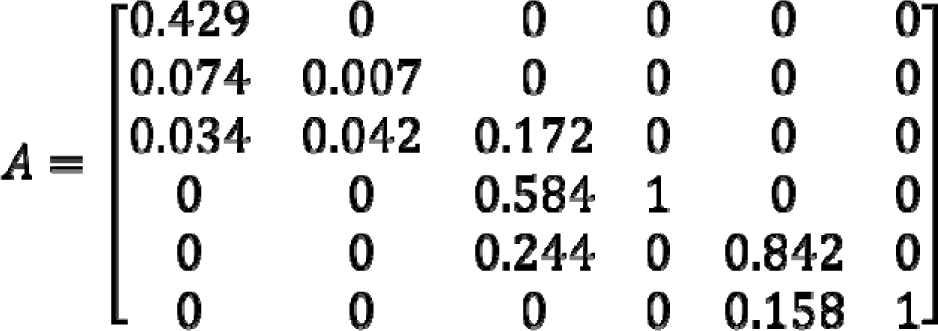

Where the diagonal corresponds to the probability of remaining in the same class after one *dt,* the subdiagonal corresponds to the probability of growing to the next class, and the subsubdiagonal the probability of jumping to the class after the next one.

**Table 1.** Criteria followed to establish the different stages of the pelagic phase of medusae based on size (from Bordehore et al. 2015). *DBW = Diagonal bell width.

| Stage | Size | Sexual and morphological characteristics |
| --- | --- | --- |
| 1 Juvenile 1 (J1) | $0.2 \text{ mm} \leq \text{DBW}^* \leq 5 \text{ mm}$ | From polyp metamorphosis until the jellyfish develops gastric cirri and pedalia. |
| 2 Juvenile 2 (J2) | $5 \text{ mm} \leq \text{DBW} \leq 10 \text{ mm}$ | Developed velarium. |
| 3 Juvenile 3 (J3) | $10 \text{ mm} \leq \text{DBW} \leq 15 \text{ mm}$ | Adult morphology without developed gonads. |
| 4 Adult 1 (A1) | $15 \text{ mm} \leq \text{DBW} \leq 20 \text{ mm}$ | With gonads. |
| 5 Adult 2 (A2) | $20 \text{ mm} \leq \text{DBW} \leq 25 \text{ mm}$ | With gonads. |
| 6 Adult 3 (A3) | $\text{DBW} \geq 25 \text{ mm}$ | With gonads. |

**Table 2.**
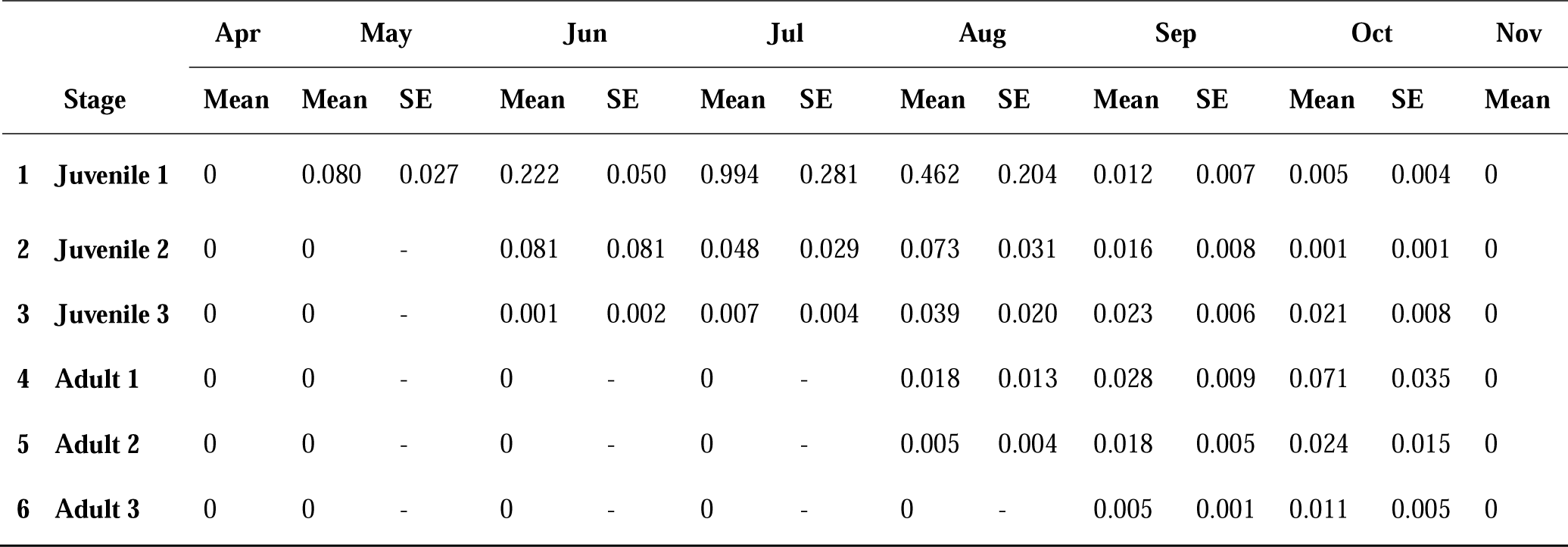
*C. marsupialis* density (medusae m^−3^ ± Standard Error of the mean, SE) by month (from Bordehore et al. 2015).

To be able to model a self-sustained and stable population, we closed the life cycle by adding into our dynamic model stocks for planulae and polyps and a strobilation pattern of polyps that generate new juveniles. Cubozoans are dioecious and oviparous, they have a bipartite life cycle consisting of a pelagic medusa phase and a sessile polyp phase which releases juvenile jellyfish through metamorphosis (polyp strobilation), without going through the ephyra phase like other cnidarians in the Scyphozoa class (Straehler-Pohl & Jarms, 2005). The life cycle begins with the mating of adults and the subsequent fertilisation of eggs. The eggs then sink to the bottom and in a few hours evolve into a planula larva that settles if an appropriate substrate is found. After a few days, the polyp develops, and one year later it can perform two types of metamorphosis (Cutress & Studebaker, 1973): the first type of metamorphosis results in the polyp transformation into medusae, through strobilation, without leaving any remnant polyp behind, occurring in the majority of cubozoan species (Toshino et al., 2015). The second type of metamorphosis, which was observed in 45% of polyps of *C. marsupialis*, results in a residue that eventually generates a small remnant polyp till the next year (Straehler-Pohl & Jarms, 2005). The life cycle is completed when the juvenile jellyfish grows to its adult size, reaches maturity, and reproduces. *C. marsupialis* life cycle exhibits a temporal seasonality in the south-western Mediterranean, appearing in juvenile sizes (0.2-5 mm) from May to October, and adult sizes (>15 mm) from August until the end of October (Bordehore et al., 2015) (see Table 2).

### Model calibration and goodness of fit

In order to create a reliable model it is necessary to calibrate it. We used two requirements to ensure that the model is correctly balanced. First, a stable population must be established over a relatively long period of time, for example 300 months (25 years). Note that the time unit, *dt*, was set at 1 month. Second, the model output must be able to replicate the observed intra-annual variation of each size class within a 95% confidence interval, estimated from the SE in Table 2.

Currently, little is known about the abundances of the benthic phase of *C. marsupialis*. Hence, we will estimate planula and polyp data based on literature and expert opinions, as well as consistency with the model itself.

Once we had considered all the biological criteria, we calibrated a time-variable strobilation pattern and the initial number of the polyp stock in order to get the optimal fit between the dynamic model output and the *in situ* measurements. The dynamic model is formed by a system of differential equations (DE). However, it is discretised into a system of finite differential equations (FDE) because the *in situ* measurements are obtained monthly (see Supplementary Material 4). This is the reason why *dt* was set to 1 month.

Then, the goodness of fit between the two models, the published matrix approach (Bordehore et al., 2015) and this dynamic model, is calculated by the root mean square (RMS) of the residual between the observed data and the model outputs. The lower the RMS value, the better the goodness of fit.

### Sensitivity analysis

A sensitivity analysis allows us to run a model multiple times to analyse the range of the output behaviour that can be generated under different initial conditions and parameter values, including parameters that follow a given probability distribution.

To understand how the different parameters affect the jellyfish population, we conducted a sensitivity analysis on the most relevant input parameters in the model: *growth rates* (with the exception of the *growth rate* from *A1* to *A2*, whose value is 0)*, settlement rate, fecundities, J1* and *J2 death rates of medusa phase* - also including *predation, advection,* and *emigration* - (the rest of stages’ *death rates* were not considered because their value is 0)*, egg to planula rate, planula settlement rate, planula death rate, polyp strobilation rate* and *strobilation pattern*, and *new remnant polyps rate*). An increase and decrease of 2, 5, 10 and 15% was applied to each non-zero parameter, one at a time, and the corresponding simulations were analysed. Nine simulations (the original plus the 8 variations) were performed for each of the 21 parameters/converters, which makes a total of 189 simulations.

Therefore, the parameters analysed were classified as robust, sensitive or very sensitive depending of how they affect to the production of J1, and according to the following qualitative criteria: if the output of the associated simulation is very similar to the 25-year base stable simulation, we say that the parameter is robust; if the output is notably different, showing long term changes that lead to a different equilibrium in the population, we will say that the parameter is sensitive; if the output produces an abrupt change in the population, the parameter is very sensitive.

### Scenarios comparison

Once the model had been calibrated to get a stable population over 25 years, we simulated different scenarios, that is, we introduced some changes in the stable population to study the evolution of the population from that point. The purpose is to learn how changes in some parameters (Figure 1) will affect the future of an initially interannual stable population. We are in search of answers to questions such as *What would happen if…?* We performed 4 scenarios:

- Scenario 1: What would happen if there is a reduction in prey availability? To answer this, we tested a decline in *growth rates* (25 and 50% reduction) reflecting a reduction in prey availability (parameters used are detailed in Table 3).
- Scenario 2: What would happen if there were different jellyfish removal strategies? To do that, we tested three strategies: (1) a 10% juvenile size jellyfish removal from June to September; (2) a 10% adult size jellyfish removal from June to September; and (3) a 10% total jellyfish removal from June to September (parameters used are detailed in Table 4).
- Scenario 3: What would happen if there is a change in the advection rate? To study this, we increased and decreased the *advection rate* by 15, 25 and 50%, respectively, to simulate a change in advection by currents on juveniles, e.g. construction or removal of breakwaters (parameters used are detailed in Table 3).
- Scenario 4: What would happen if there are changes in substrate availability? We tested these changes varying the success of planula larvae settlement by 5, 10 and 20% (parameters used are detailed in Table 3).

**Figure 1.**
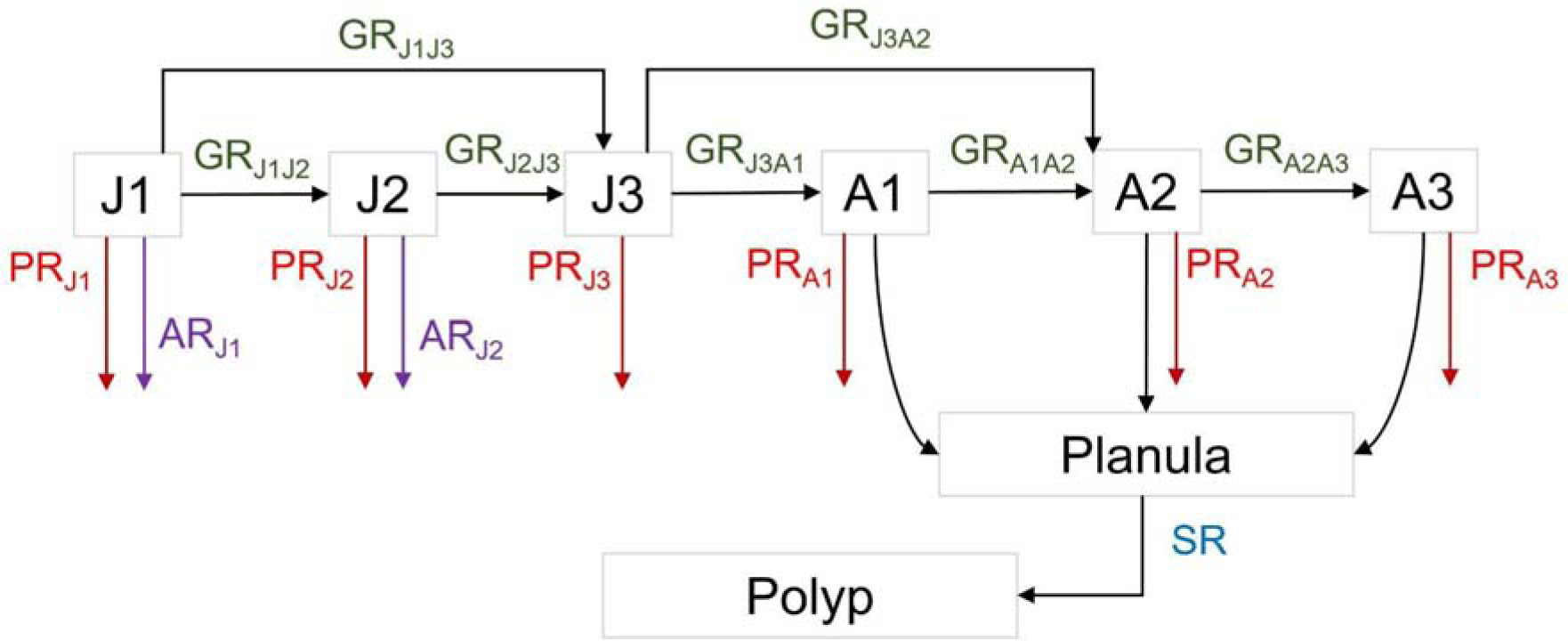
Simplified diagram representing the parameters of biological interest used in the scenarios. GR=Growth rate, PR=Predation rate, AR=Advection rate and SR=Settlement rate.

**Table 3.** Decline in prey availability (scenario 1), change of advection rates (scenario 3) and changes in settlement rate (scenario 4) scenarios carried out for *C. marsupialis*. References for each scenario are Bordehore et al. 2015**^1^**, Bordehore et al. 2019**^2^** and Boero, 2013**^3^**. GR=Growth rate, PR=Predation rate, AR=Advection rate and SR=Settlement rate.

| | Rate | $GR_{J1J2}^1$ | $GR_{J1J3}^1$ | $GR_{J2J3}^1$ | $GR_{J3A1}^1$ | $GR_{J3A2}^1$ | $GR_{A1A2}^1$ | $GR_{A2A3}^1$ | $PR_{J1}^1$ | $PR_{J2}^1$ | $AR_{A1}^{2,3}$ | $AR_{A2}^{2,3}$ | $SR^3$ |
| --- | --- | --- | --- | --- | --- | --- | --- | --- | --- | --- | --- | --- | --- |
| Scenario | Base | 0.074 | 0.034 | 0.042 | 0.584 | 0.244 | 0 | 0.158 | 0.155 | 0.317 | 0.154 | 0.317 | 0.020 |
| 1 | -25% GR | 0.056 | 0.026 | 0.032 | 0.438 | 0.183 | 0 | 0.118 | - | - | - | - | - |
|  | -50% GR | 0.037 | 0.017 | 0.021 | 0.292 | 0.122 | 0 | 0.079 | - | - | - | - | - |
| 3 | -50% AR | - | - | - | - | - | - | - | - | - | 0.077 | 0.159 | - |
|  | -25% AR | - | - | - | - | - | - | - | - | - | 0.116 | 0.238 | - |
|  | -15% AR | - | - | - | - | - | - | - | - | - | 0.131 | 0.269 | - |
|  | +15% AR | - | - | - | - | - | - | - | - | - | 0.177 | 0.365 | - |
|  | +25% AR | - | - | - | - | - | - | - | - | - | 0.193 | 0.396 | - |
|  | +50% AR | - | - | - | - | - | - | - | - | - | 0.231 | 0.476 | - |
| 4 | -15% SR | - | - | - | - | - | - | - | - | - | - | - | 0.017 |
|  | -10% SR | - | - | - | - | - | - | - | - | - | - | - | 0.018 |
|  | -5% SR | - | - | - | - | - | - | - | - | - | - | - | 0.019 |
|  | +5% SR | - | - | - | - | - | - | - | - | - | - | - | 0.021 |
|  | +10% SR | - | - | - | - | - | - | - | - | - | - | - | 0.022 |
|  | +15% SR | - | - | - | - | - | - | - | - | - | - | - | 0.023 |

**Table 4.**
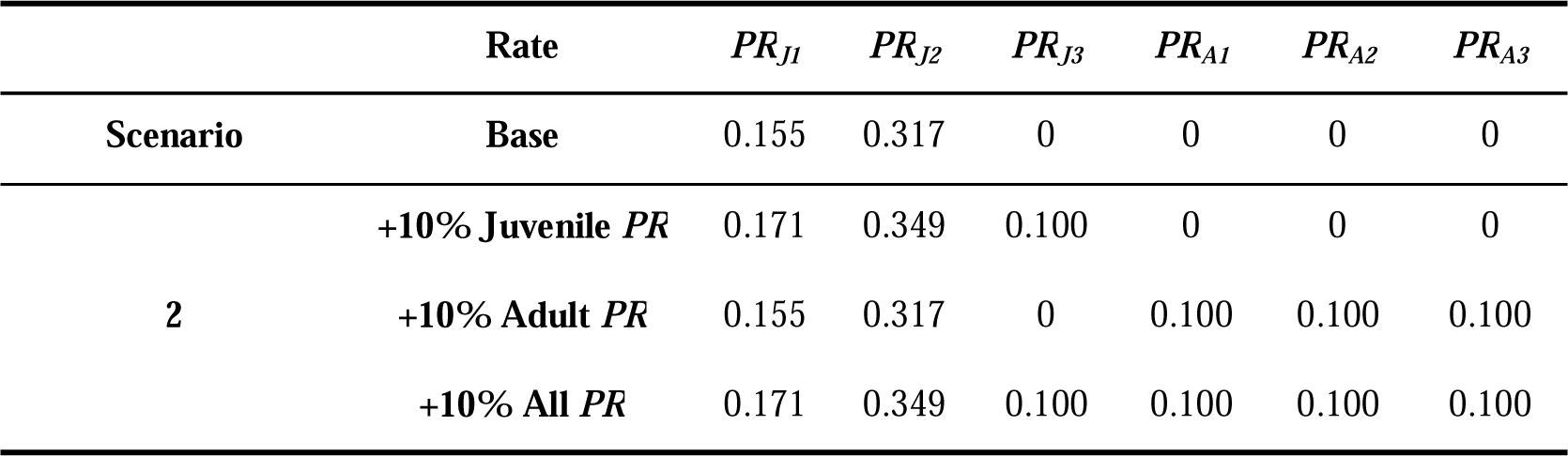
Jellyfish removal scenarios (scenario 2) carried out for *C. marsupialis*. References for each scenario are Bordehore et al. 2015. PR=Predation rate.

| | Rate | $PR_{J1}$ | $PR_{J2}$ | $PR_{J3}$ | $PR_{A1}$ | $PR_{A2}$ | $PR_{A3}$ |
| --- | --- | --- | --- | --- | --- | --- | --- |
| Scenario | Base | 0.155 | 0.317 | 0 | 0 | 0 | 0 |
| 2 | +10% Juvenile PR | 0.171 | 0.349 | 0.100 | 0 | 0 | 0 |
|  | +10% Adult PR | 0.155 | 0.317 | 0 | 0.100 | 0.100 | 0.100 |
|  | +10% All PR | 0.171 | 0.349 | 0.100 | 0.100 | 0.100 | 0.100 |

## RESULTS

### Dynamic model for *C. marsupialis*

The dynamic model we developed is shown in Figure 2 and more details are available in Supplementary Material 1 (model output), Supplementary Material 2 (script of the model) and Supplementary Material 3 (model in STELLA Architect format with .stmx file). Note that this model includes both the pelagic and benthic phases while in the original matrix model (Bordehore et al. 2015) only the pelagic (medusa) phase was included in the transition matrix values. When an individual does not grow or stay in a size class after a *dt*, it means that it has disappeared from our studied population, which could be attributed to different biological causes, *i.e.* predation, emigration (actively exits the population by swimming), advection (passively exits the population due to currents) or death. Although we cannot evaluate the contribution of these four causes, we consider them relevant in order to perform different scenarios, to be able to split these *death* causes. The benthic phase is formed by the stocks of planula larvae and polyps (including remnant polyps from the previous year) and processes such as metamorphosis from polyps, planula larvae settlement, or death.

**Figure 2.**
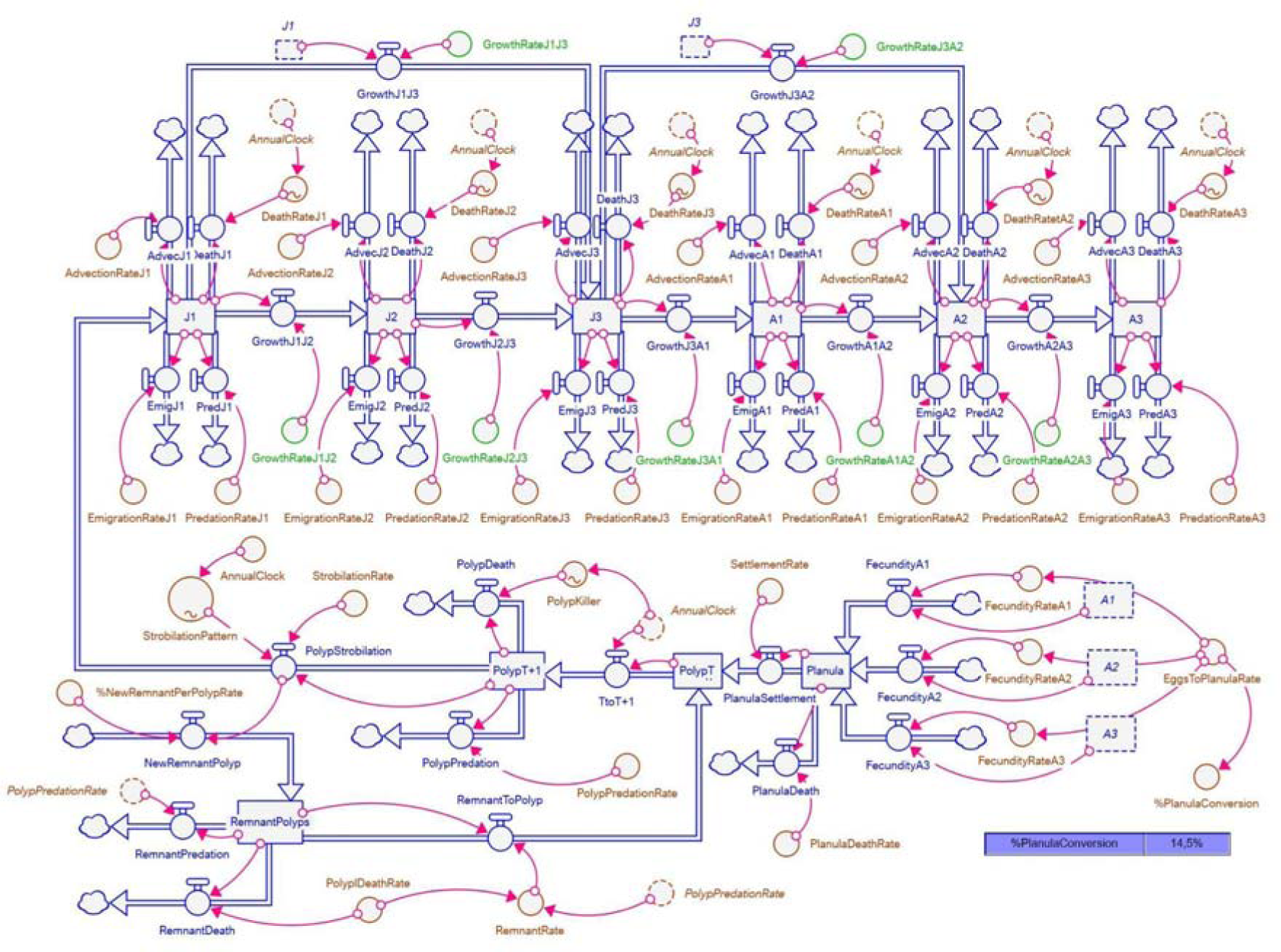
Dynamic model of *C. marsupialis* in STELLA Architect. The entire model script can be downloaded in Supplementary Material 2, the STELLA Architect .stmx file can be downloaded from Supplementary Material 3 and the associated finite differential equations (FDE) and calibration procedures are described in Supplementary Material 4.

### Model calibration and goodness of fit

Based on the use of the values of the transition matrix (Equation 1), and in order to calibrate the model, we needed to estimate or calculate the values of the rest of the parameters. The aim was to obtain a model that simulates: (1) the intra-annual population for each size class as close as possible to the observed data; (2) a stable population for a long period, e.g. 25 years (300 months). The values of the parameters were obtained in two different ways, either from the literature or by deducing them from the dynamic model.

From the literature we obtained: (1) the probabilities of stay or growth of each medusa stage were obtained from the transition matrix (Bordehore et al., 2015) and can be found in Table S1 of Supplementary Material 4; (2) in the absence of specific data for *C. marsupialis*, and considering the similarity of both species, we used the data of fecundity and planulae production from *Morbakka virulenta* (Kishinouye, 1910) from Thosino et al., (2013); (3) the percentage of polyps that strobilate each year and the percentage that remains were obtained from Straehler-Pohl & Jarms (2005).

From the dynamic model, we calculated the total annual population of polyps and the discrete probability density function of the monthly polyps strobilation (hereafter *strobilation pattern*). Note that the polyps population does not refer to the real number of polyps, but a number that represents all the actual polyps and that can be altered to test proportional changes. The strobilation pattern is one of the cornerstones of the model and represents the relative number of polyps strobilating each month. The values of the strobilation pattern can be found in Table S2 of Supplementary Material 4. Further details on the model calibration can be found in Supplementary Material 4.

The calibrated model produces an abundance output that is within the 95% confidence interval of the field data for each month of each stock, except for September and October of *J1* and *J2*, and September of *A3* (Figure 3). This means that there are only 5 months of disagreement. On the contrary, the output of the matrix model had 18 months of disagreement within the 95% confidence interval (*J1*: May, June, September and October; *J2*: October; *J3*: June, July, November and December; *A1*: June, September and December; *A2*: June, November and December; *A3*: July, September and December).

**Figure 3.**
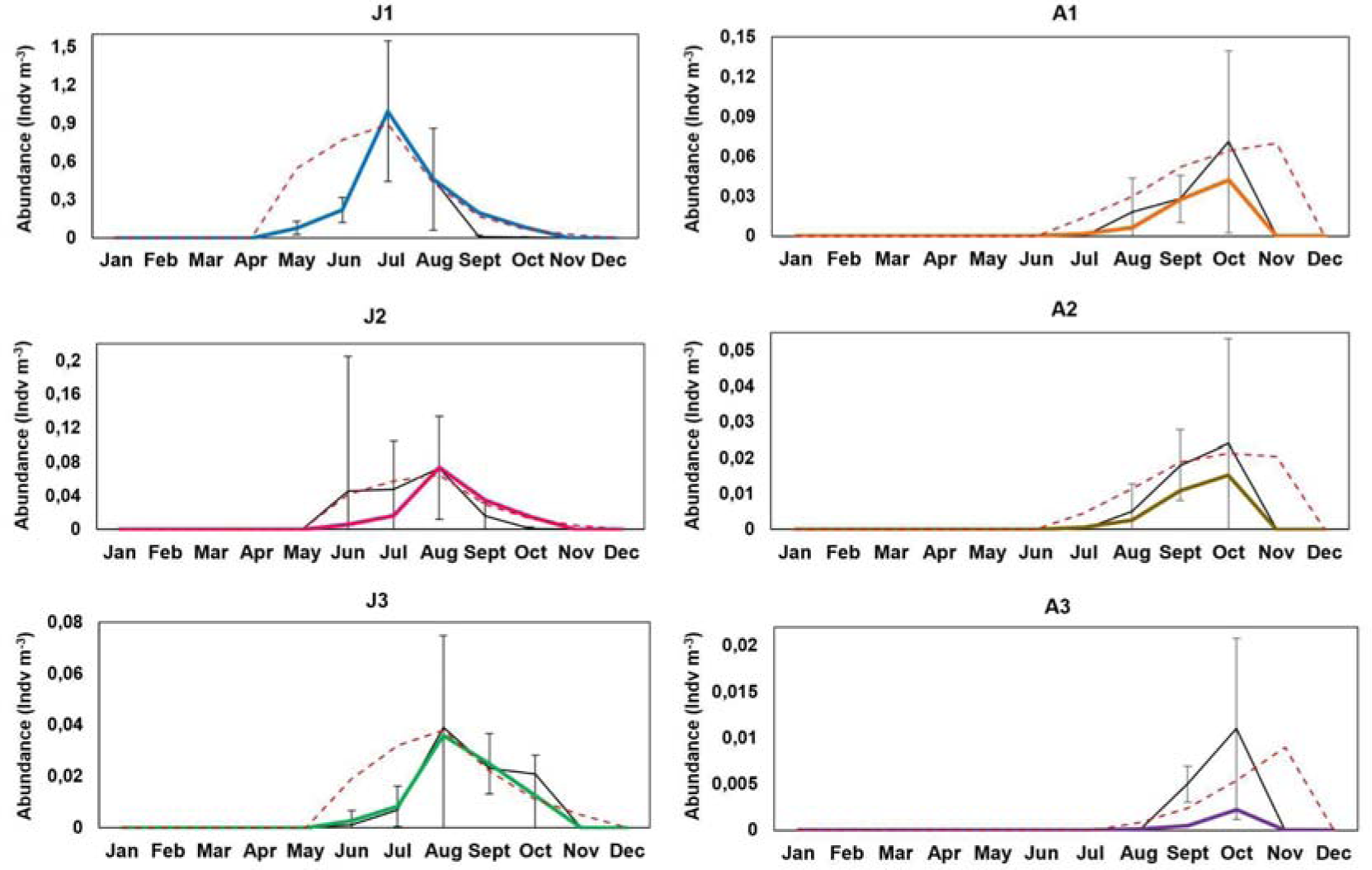
Abundances of *Carybdea Marsupialis* from in situ measurements from Bordehore et al. 2015, including the 95% confidence intervals (black lines), matrix model (dashed red line), and the dynamic model in Figure 2 (coloured lines).

The RMS of the residual between the observed data and the output of the models, both matrix and dynamic, quantifies the goodness of fit. The RMS has been calculated individually for each jellyfish stage (Table 5). Except for the *J2* stage, all stages had a lower RMS in the dynamic model than in the matrix model, and therefore a better fit in the former. If we calculate the RMS of all stages at the same time, we get 0.0065 for the dynamic model and 0.0229 for the matrix model. Note that the RMS of the residuals was calculated only for months where the data, the model output or both were non-zero. In addition, the calibrated model was able to produce a stable population over 25 years (Figure 4).

**Figure 4.**
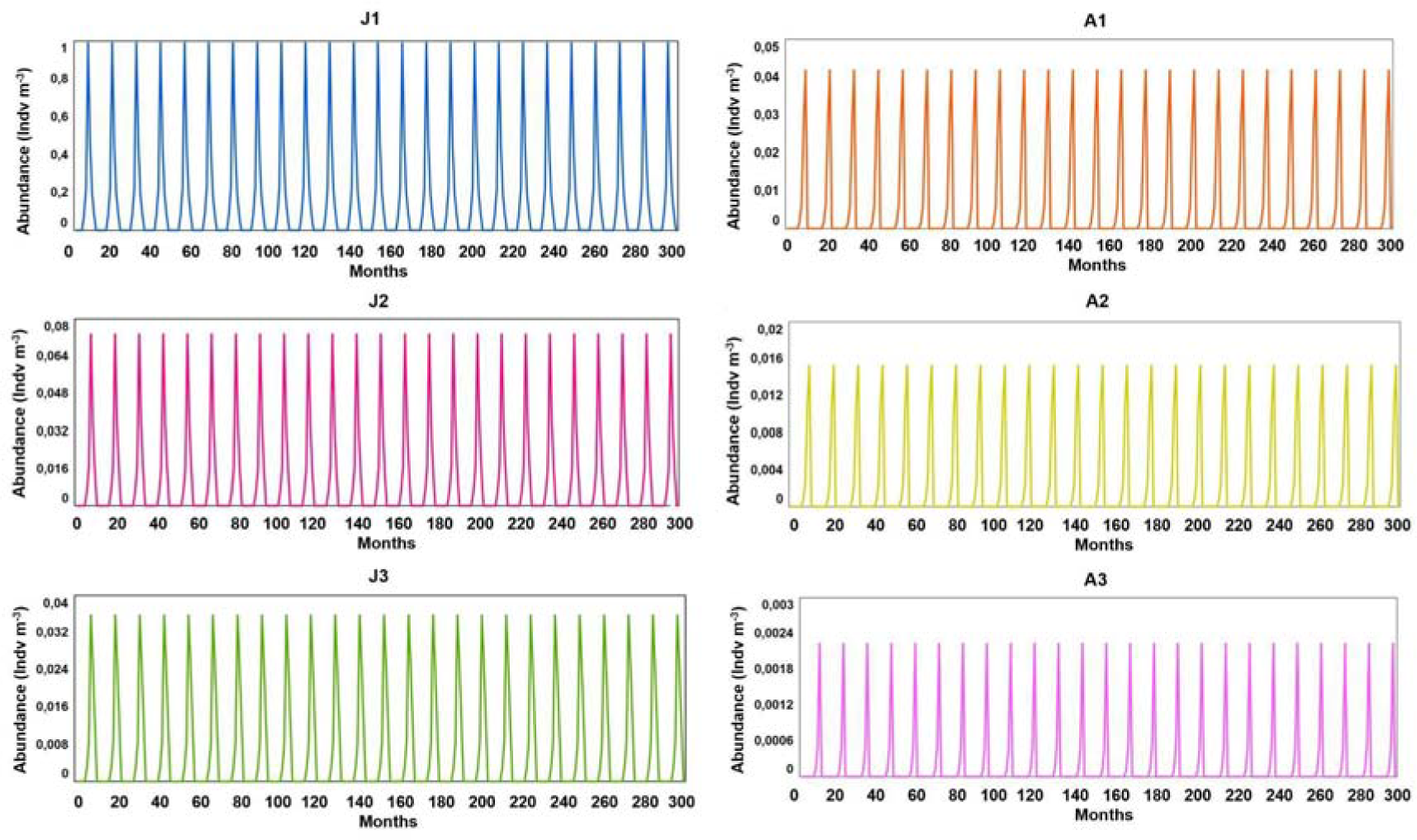
Output (abundances of each size-class, individuals m^−3^) of the calibrated dynamic model of *C. marsupialis* over 300 months (25 years). Note that each maximum corresponds to 1 year, the one represented in Figure 3 for each size-class.

**Table 5.**
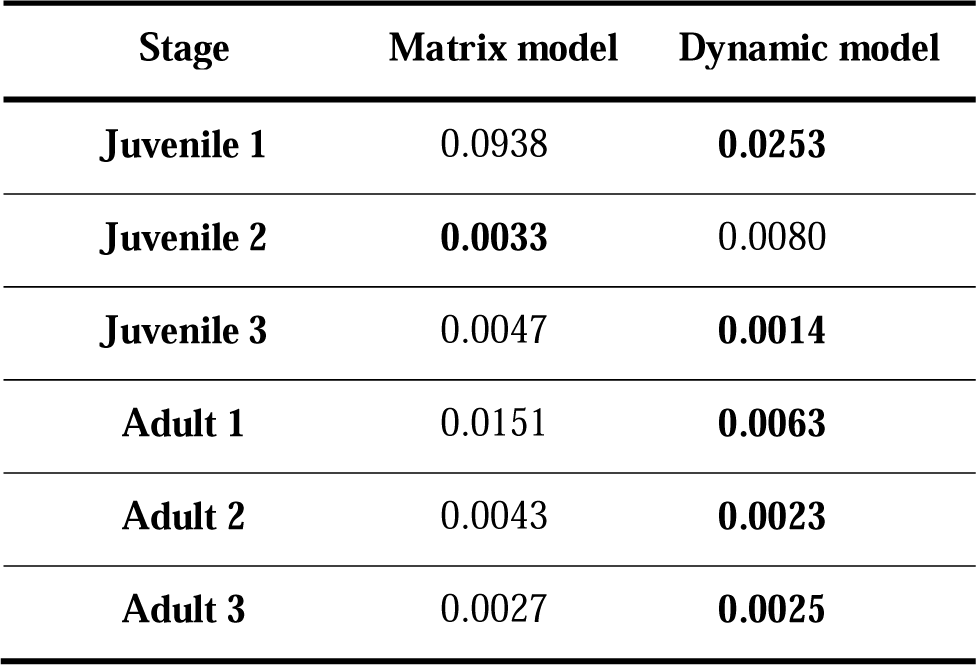
RMS (root mean square) of the residual between the observed data and the output of the matrix (from Bordehore et al., 2015) and dynamic models (this study). The RMS was calculated only for months where the field data, the model’s output or both were non-zero. Better adjustment marked in bold.

### Sensitivity analysis

We conducted a sensitivity analysis for those parameters that were of biological interest in the model in order to assess the influence on the model output. The analysis consisted in which was measured through the effect on *J1* production per year. We show here the main results (Table 6) while a complete report can be found in Supplementary Material 5.

**Table 6.**
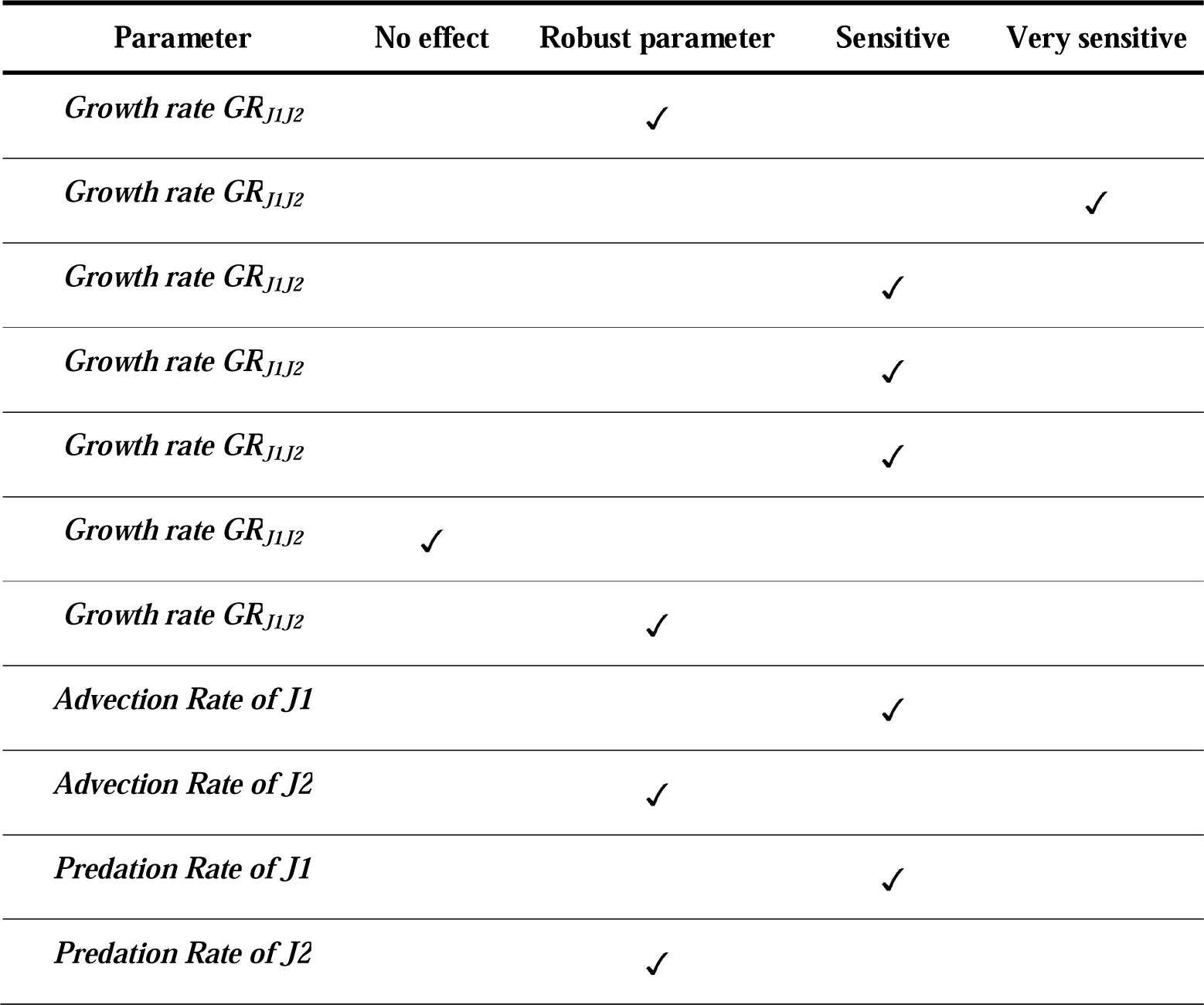
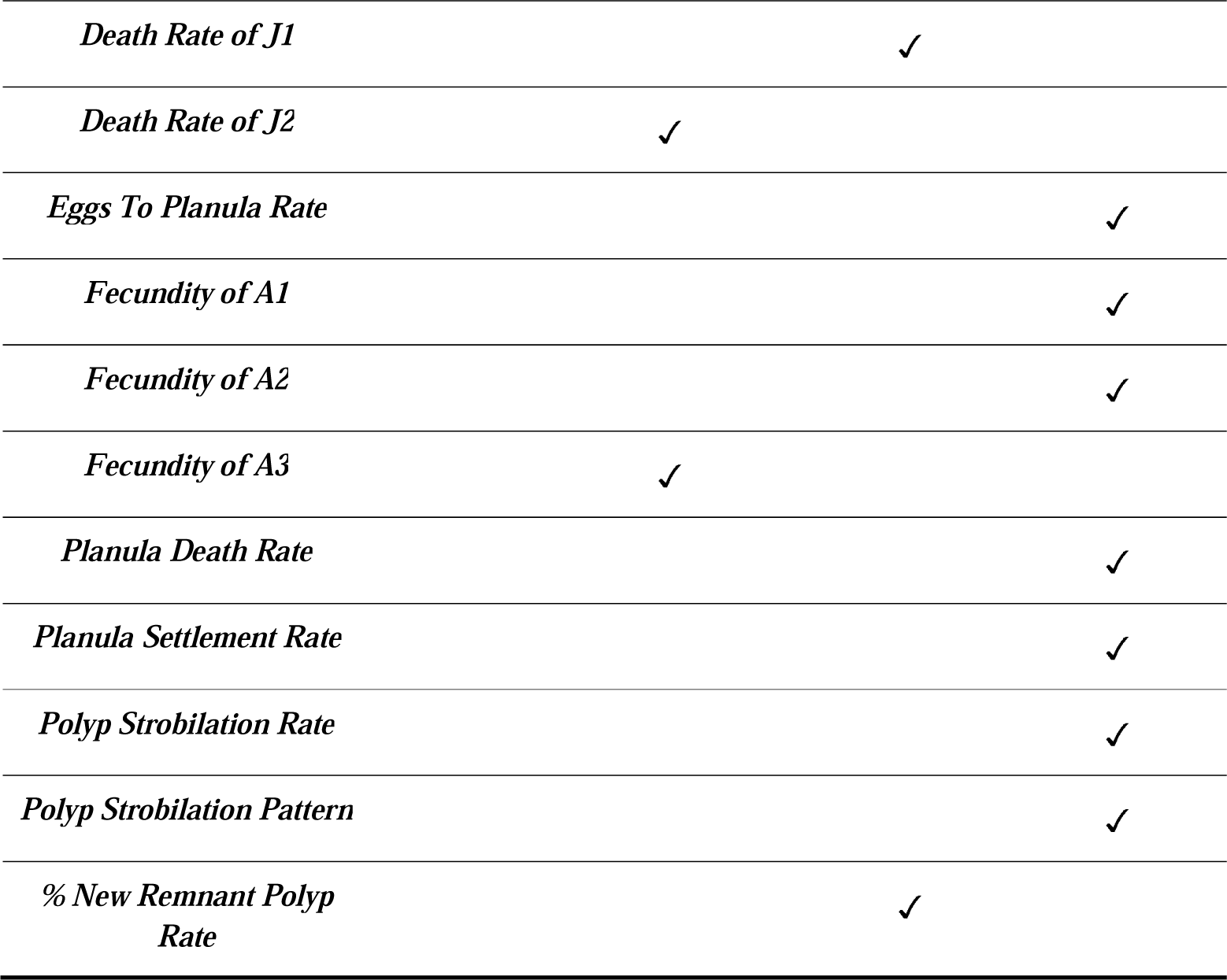
Results of the sensitivity analysis. Note that the reference output we looked at was the *J1* individuals production per year.

The sensitivity analysis for *growth rate* exhibited different effects on jellyfish abundances depending on which stage rate was modified. *GR_A1A2_* was not analysed because its value was 0. *GR_J1J2_* and *GR_A2A3_* were robust, since all the stock barely showed changes in the 25 years after the increases and decreases in the variables. *GR_J2J3_*, *GR_J3A1_* and *GR_J3A2_* were sensitive to changes but they did not have a big impact. Finally, *GR_J1J3_* was the most sensitive of the *growth rates*, showing the largest deviation from the 25-year base simulation. If *GR_J1J3_* is decreased (increased) by 2%, the *J1* production after 25 years will decrease (increase) by 25% (42%).

Variations in *death, predation* and *advection* of *J1* produced almost identical outputs. In all cases, the three variables were quite robust, showing changes in the *J1* production lower than 25% after 25 years. For the *death*, *predation* and *advection rates* of *J2*, the same almost identical outputs were displayed for each simulation of the analysis, showing that the three were very robust.

The *fecundity* of *A1* was very sensitive. A small increase (decrease) of 2% produced a large increase (decrease) of *J1* production of 25% (33.33%) after 25 years. A moderate increase (decrease) of 15% produced a large increase (decrease) of *J1* production of 433% (83%). *A2* fecundity behaved quite similarly but with lower sensitivity: an increase (decrease) of 2% produced an increase (decrease) of 8% (8%), while an increase (decrease) of 15% produced an increase (decrease) of 125% (58%). The analysis of *A3* was very different, whose fecundity was very robust.

In the case of the planula *death rate*, we found another very sensitive parameter. A decrease of 2% caused an increase of the *J1* production of 42%, while an increase of 2% produced a decrease of 8%. If the planula *death rate* is decreased (increased) by 15%, the *J1* production increases (decreases) 1567% (75%).

The planula *settlement rate* was also very sensitive. A 2% increase led to a rise of the output of 42%, while a 2% reduction caused a 33% drop. When the increase reached 15%, the *J1* production increased by 1150%, while a 15% reduction produced a decrease of 93%.

The *% New Remnant polyp rate* was a highly sensitive parameter since an increase of 15% resulted in 117% increase in *J1* production, and a decrease of 15% resulted in 50% decrease. However, this parameter is not as sensitive as both polyp *strobilation rate* and *strobilation pattern*. When the latter increased (decreased) by 2%, the *J1* production increased (decreased) by 50% (33%). These two strobilation variables were the most sensitive of the entire model because an increase (decrease) of 15% increased (decreased) the *J1* production by 1900% (98%). Further details can be found in Supplementary Material 5.

### Scenarios simulation

#### Scenario 1: Decline in prey availability

Scenarios dealing with prey availability were simulated by reducing the same percentage of all the model’s *growth rates* at the same time. Under a 25% reduction, the abundance decreased 75% in 5 years, 96% in 10 years and reached values close to zero at year 21 (month 247; Figure 5). Under a 50% reduction, the abundance decreased 95% in 5 years and reached values close to zero by the middle of year 10 (month 155).

**Figure 5.**
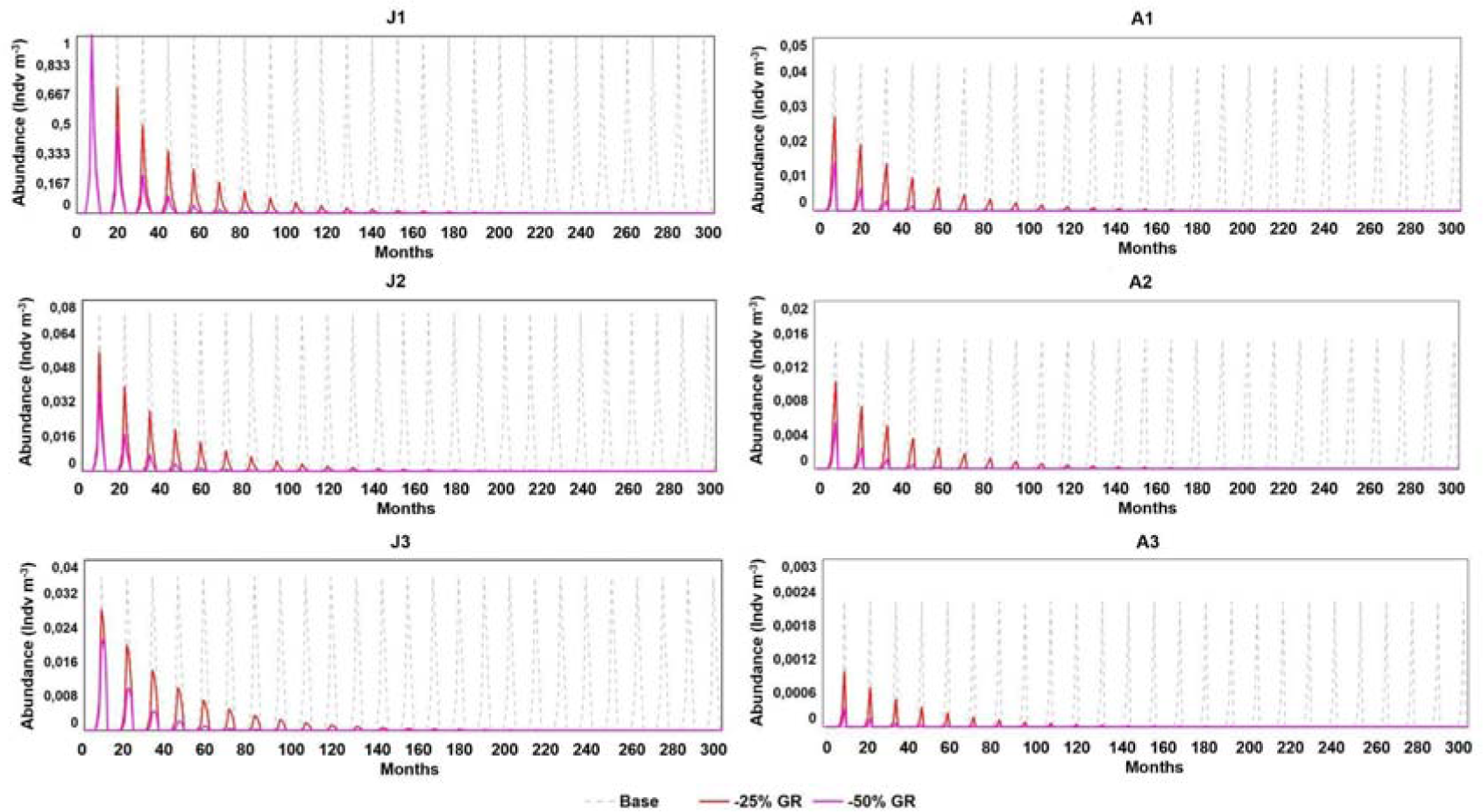
Simulations of Scenario 1: Decline in prey availability scenarios compared with the base model simulation. GR (Growth rate). We compared the base scenario (dashed grey), a decrease of 25% in growth rate (red line) and 50% (pink line). *J1*: juvenile stage 1; *J2*: juvenile stage 2; *J3*: juvenile stage 3; *A1*: adult stage 1; *A2*: adult stage 2; *A3*: adult stage 3.

#### Scenario 2: Jellyfish removal strategies

This scenario envisages the implementation of jellyfish removal strategies. The outputs produced in the jellyfish removal scenarios showed that when we removed a monthly 10% of juvenile population (*J1*+*J2*+*J3*) from June to September every year, the total jellyfish abundance decreased 16% in 5 years, 32% in 10 years and 64% in 25 years. When we removed 10% of the adult population (*A1*+*A2*+*A3*) during the same months, the total jellyfish abundance decreased 16% in 5 years, 33% in 10 years and 66% in 25 years. Finally, if we removed 10% of the total jellyfish, abundance decreased 30% in 5 years, 60% in 10 years and 88% in 25 years (Figure 6).

**Figure 6.**
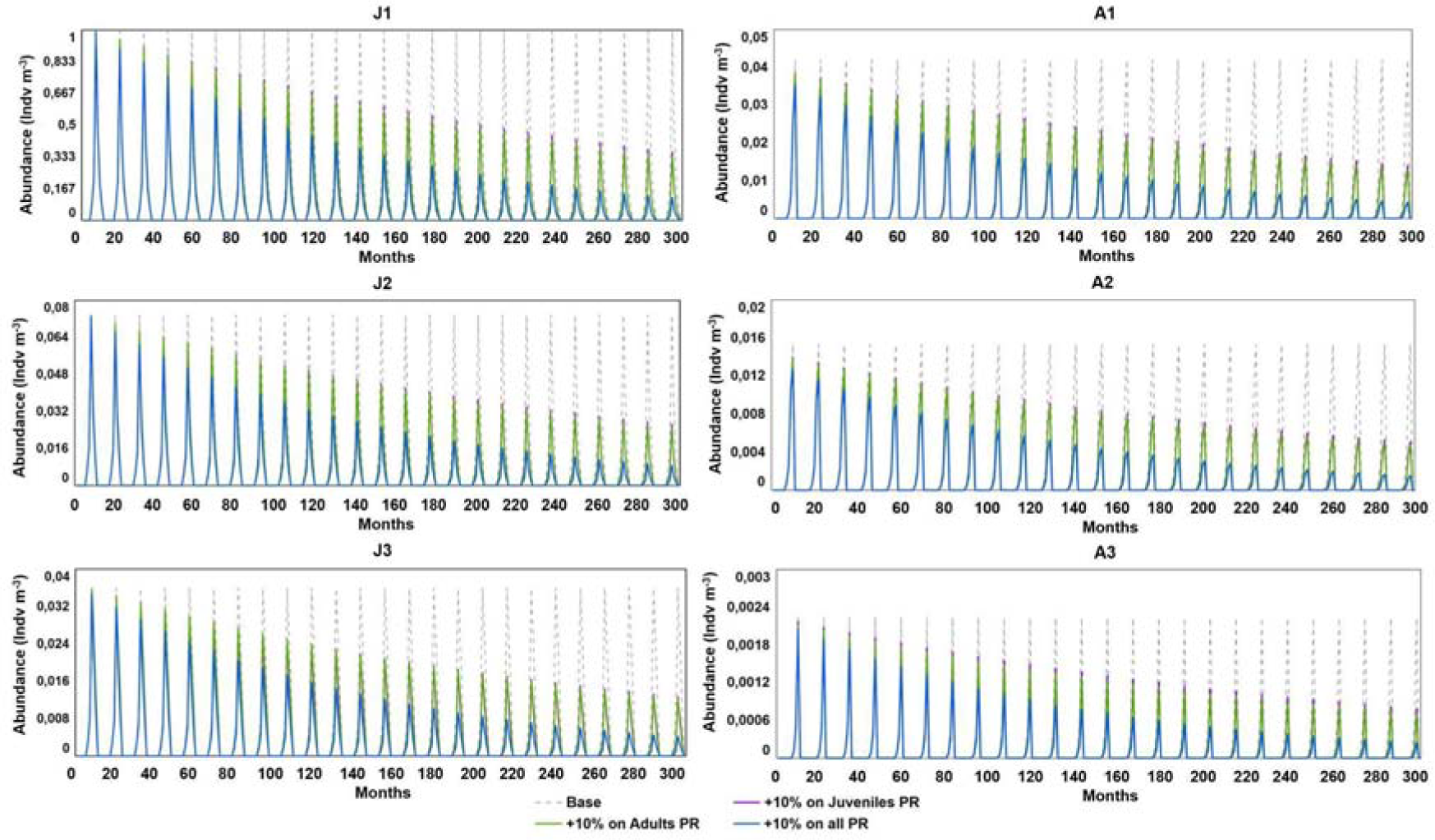
Simulations of Scenario 2: Jellyfish removal scenarios comparison. PR (Predation rate). We compared the base scenario (dashed grey line), only juvenile jellyfish removal (purple), only adult removal (green) and all jellyfish sizes removal (blue). *J1*: juvenile stage 1; *J2*: juvenile stage 2; *J3*: juvenile stage 3; *A1*: adult stage 1; *A2*: adult stage 2; *A3*: adult stage 3.

#### Scenario 3: Change of advection rates

A 15%, 25% and 50% decrease in advection rate led to an increase of 4%, 8% and 16% respectively of the jellyfish abundance after 5 years, 9%, 16% and 42% respectively after 10 years, and 26%, 47% and 117% respectively after 25 years. By contrast, an increase of advection rate by 15%, 25% and 50% led to a population decrease of 4%, 7% and 13% respectively of the population after 5 years, 9%, 14% and 26% respectively after 10 years and 16%, 32% and 54% respectively after 25 years (Figure 7).

**Figure 7.**
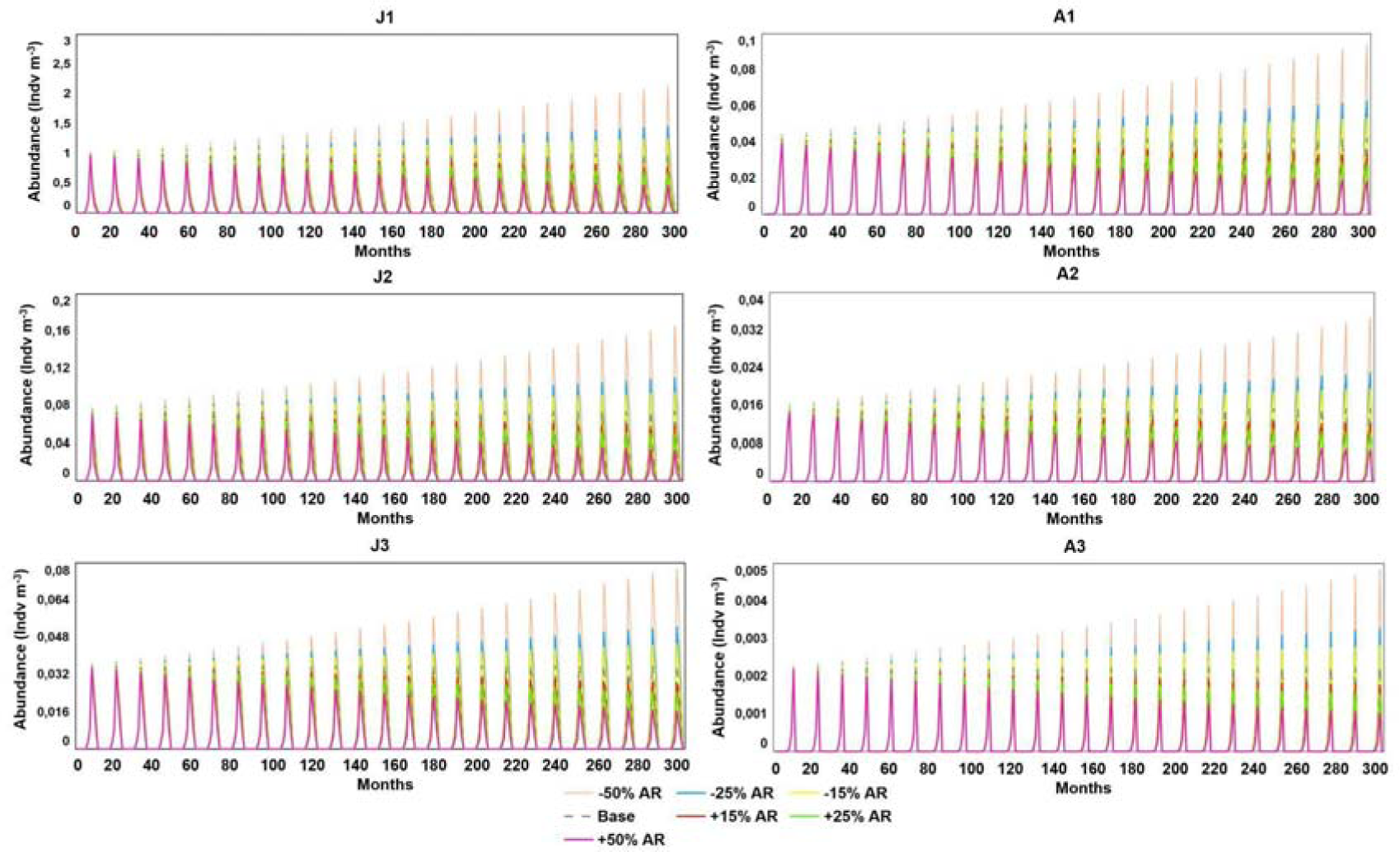
Simulations of Scenario 3: Advection rate scenarios compared with the base model simulation. AR (Advection rate). We compared the base scenario (dashed grey line) with six scenarios: a decrease in AR of 50% (beige line), 25% (blue line), and 15% (yellow line) and an increase of AR of 15% (red line), 25% (green line) and 50% (magenta line). *J1*: juvenile stage 1; *J2*: juvenile stage 2; *J3*: juvenile stage 3; *A1*: adult stage 1; *A2*: adult stage 2; *A3*: adult stage 3.

#### Scenario 4: Changes in settlement rate

Finally, *the settlement rate* (SR) scenario simulated changes in substrate availability. For a reduction of 5%, 10% and 15% of the SR the jellyfish abundance decreased by 14%, 27% and 38% respectively in 5 years; 29%, 51% and 66% respectively in 10 years; and 61%, 85% and 95% respectively in 25 years (Figure 8).

**Figure 8.**
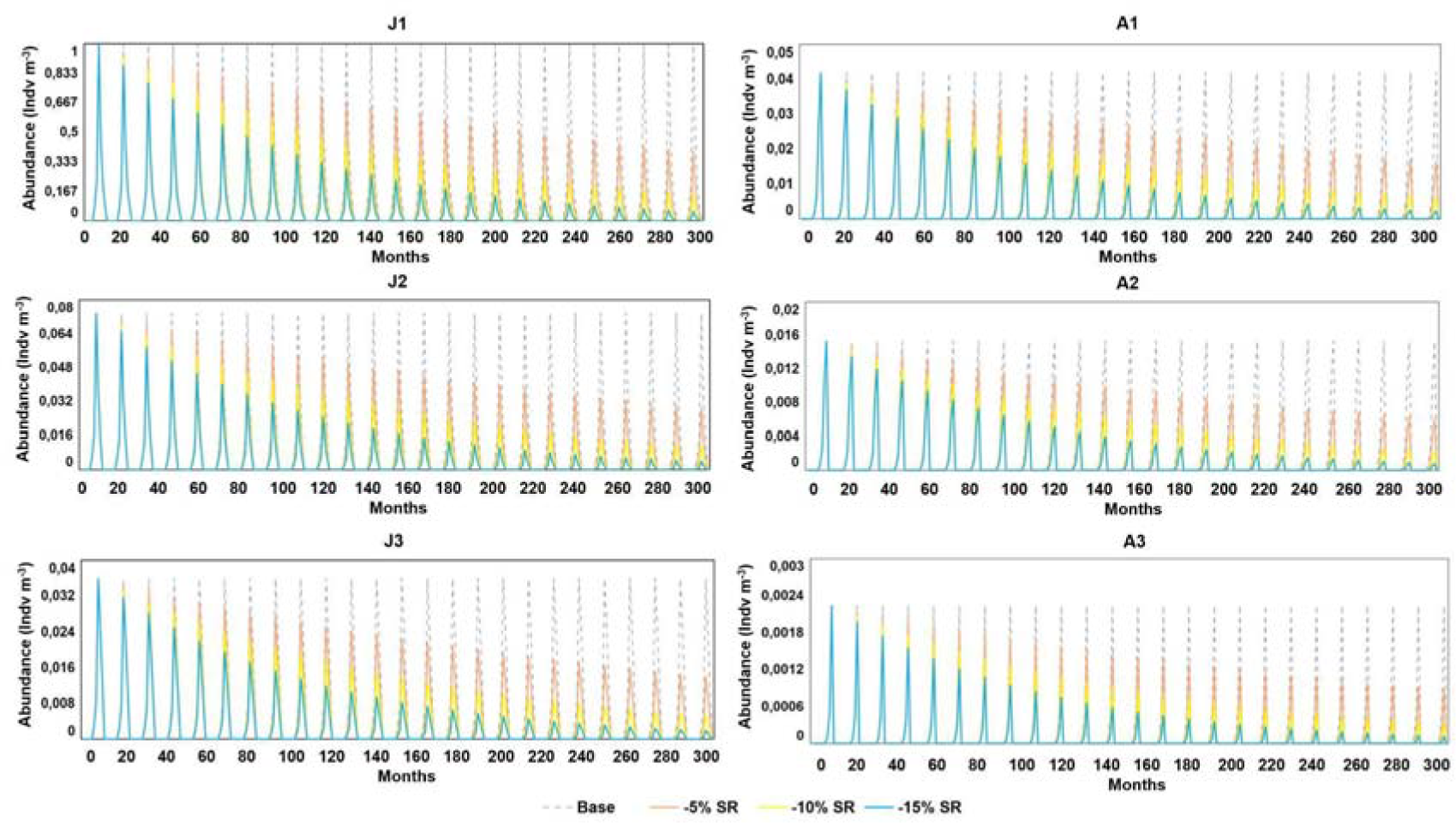
Scenario 4: Settlement rate (SR) decreasing change scenarios, simulating changes in substrate availability. We compared the base scenario (dashed grey) with a decrease of SR of 5% (beige), 10% (yellow) and 15% (blue). *J1*: juvenile stage 1; *J2*: juvenile stage 2; *J3*: juvenile stage 3; *A1*: adult stage 1; *A2*: adult stage 2; *A3*: adult stage 3.

An increase of SR of 5%, 10% and 15% led to an increase of the population’s abundance of 16%, 34% and 55% respectively in 5 years; 40%, 95% and 166% respectively in 10 years; and 147%, 490% and 1264% respectively in 25 years (Figure 9).

**Figure 9.**
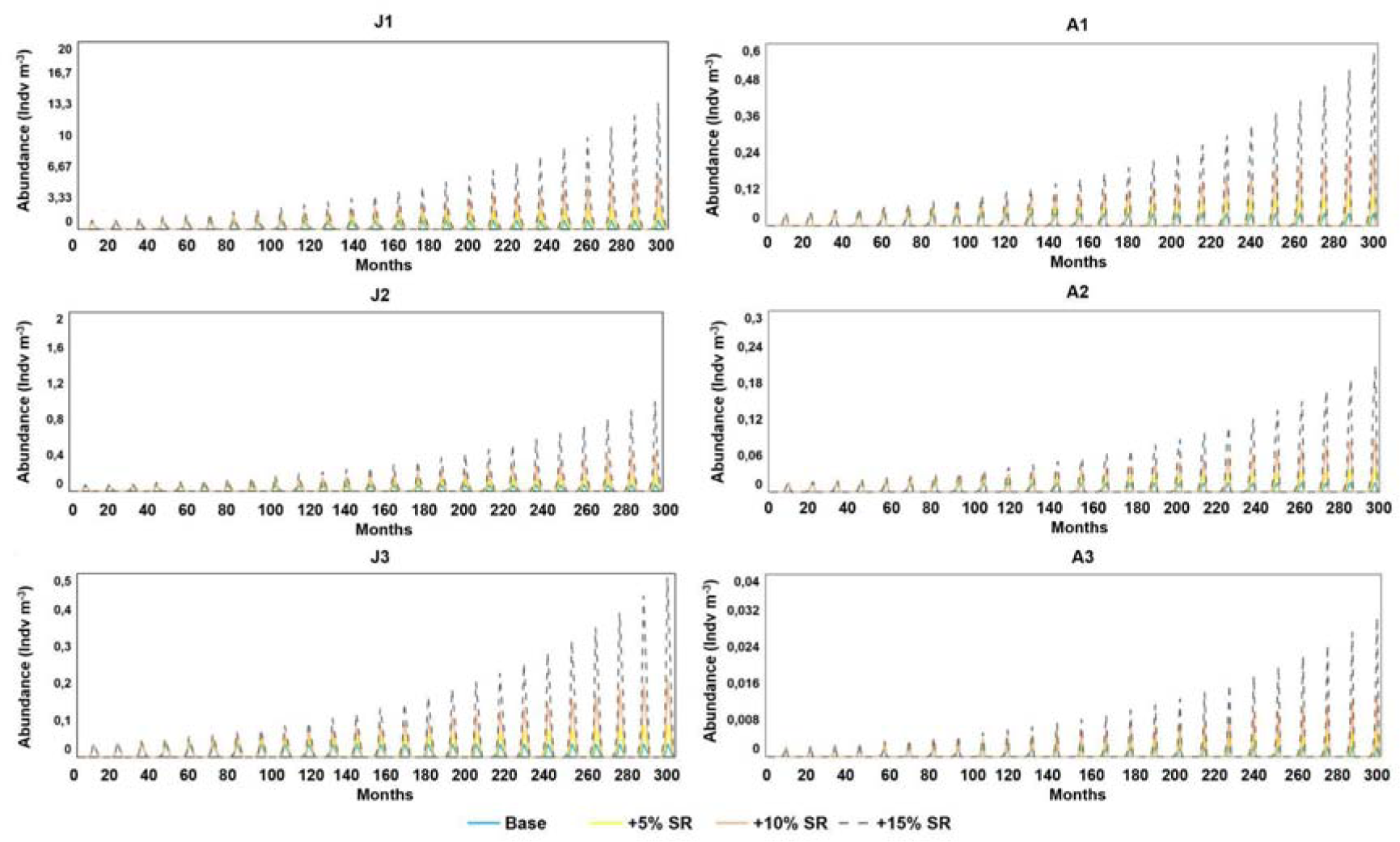
Scenario 4: Settlement rate (SR) increasing change scenarios, simulating changes in substrate availability. We compared the base scenario (blue) with an increase of SR of 5% (beige), 10% (yellow) and 15% (dashed grey). *J1*: juvenile stage 1; *J2*: juvenile stage 2; *J3*: juvenile stage 3; *A1*: adult stage 1; *A2*: adult stage 2; *A3*: adult stage 3.

Further details for all scenarios are shown in Supplementary Material 6.

## DISCUSSION

### Dynamic model

The results showed that the dynamic model improved the approximation to the observed data (Table 2), with respect to the matrix model (Table 5). The dynamic model reproduced the life cycle of the species more accurately than the matrix model since the former includes the benthic phase and an optimised monthly strobilation pattern, while the latter included an emergence vector (equation 5 in Bordehore et al., 2015) which represented the initial seeding population (note that in Bordehore et al., 2015 this emergence vector fed the population during the first 4 *dt* of the simulation). In the dynamic model, the seeding was a certain number of polyps, driven by a strobilation pattern, a more biological approximation to the real world. Furthermore, the number of juveniles produced each year depends directly on the adult population of the previous year, unlike the matrix model.

However, while this dynamic model helps to obtain valuable insights into the population dynamics of a species, it is constrained by some limitations. One major challenge is the difficulty in obtaining real data for some of the parameters that appear in the model, in this case related to the benthic phase. This is because there is very quantitative data available (e.g. time series, strobilation patterns, eggs production) not only for this species but for cnidarians and particularly for cubozoans. To overcome this data scarcity, we extrapolated some values from other similar species e.g. fecundity data from japanese box jellyfish *Morbakka virulenta* (Toshino et al., 2013). The rest of the data were estimated from the known parameters, for example, the values for *advection, predation, death* and *emigration rates* of each size-class, whose sum of values plus the growth rate must equal 1 minus the stock’s permanence, that is the diagonal of the transition matrix. To split those four concepts (*advection, predation, death* and *emigration rates*), we applied biological criteria e.g. advection should reach the maximum rates at the smallest size-class and active emigration should be the highest at bigger sizes. A relevant positive aspect of the dynamic models is that even though some data is obtained from extrapolations and indirect calculations, the results of the scenarios comparison and sensitivity analysis are valid since variations are done on relative numbers.

### Model calibration

Our dynamic model fits the observed data reasonably well (Table 2), noting that almost all abundances of jellyfish sizes fell within the 95% confidence intervals of the observed data. With regard to the 95% confidence threshold, only *J1* and *J2* in September and October, and *A3* in September fell out of the confidence interval (Figure 3).

Regarding the RMS of the residuals between the models (both dynamic and matrix) and the observed data, we obtained lower values for the dynamic model globally and for its stocks individually, except for *J2*. This means that the dynamic model approximated the observed data better.

Some converters needed a better adjustment in the dynamic model to replicate as accurately as possible the dynamics of the species. Two examples were the *death rate* for every *C. marsupialis* stage and the *polyp killer* converter (see benthic phase in Figure 2). Both variables were used as auxiliary variables (with graphical functions) to kill all individuals at a certain month when experimental data showed no presence of those size-classes or to kill non-remnant polyps once they strobilate.

### Sensitivity analysis

The highest sensitivity to changes was exhibited by the parameters related to the benthic stages such as *strobilation rate, strobilation pattern*, planula *death rate* or *settlement rate*, and the lowest sensitivity by *GR_J1J2_*, *GR_A2A3_* and *J2 predation*, *advection* and *death* rates. The rest of the analysed parameters showed a moderate sensitivity. *GR_J1J3_* was the most sensitive of the *growth rates*, reflecting the importance of the acceleration of the growth (jumping from *J1* to *J3* in one *dt* that is 1 month) in the stability of a population. Note that in the original transition matrix, *GR_A1A2_* value was 0, which means that all *A2* came directly from *J3*, reflecting that a *dt* smaller than 1 month could have modelled this rapid growth in a better way, and also that the “slow” growth from *J3* to *A1* was a dead end since all individuals that entered *A1* finally die.

### Scenarios

#### Decline in prey availability

When analysing the scenario of prey reduction, reflected in a *growth rate* lowering, the simulation showed that a 25% reduction in the *growth rate* would lead to the population disappearing in 15 years and a 50% reduction in 10 years. This scenario gave similar dynamics results to those analysed in Bordehore et al. (2015). A small decrease in prey availability could significantly lower the entire jellyfish population, showing a strong bottom-up cascade effect. This simulation supports that a removal of nutrient input from agriculture and wastewater can be translated into a reduction of primary and secondary production that in turn would reduce jellyfish abundance through lowering the *growth rates*.

An example of this bottom-up effect in the opposite direction, that is increasing prey availability, can be found in the Mar Menor (Murcia, south-east Spain). There a hypersaline coastal lagoon with nitrogen and phosphorus run-off from surrounding crops induced jellyfish blooms from 1998 (Velasco et al., 2006; Pérez-Ruzafa et al., 2002) until 2016-2017, when the population disappeared due to a hyper-eutrophication that completely altered the coastal ecosystem, including hypoxia and anoxia in some parts of the lagoon. This sudden reduction in jellyfish populations was attributed to the destabilisation of the lagoon’s trophic web due to the excessive input of nutrients, specially nitrates and phosphorus, rising temperature trends and more frequent torrential rains (Fernández-Alías et al., 2022).

#### Jellyfish removal strategies

Jellyfish removal strategies, in line with Bordehore et al. (2015), emphasise the effectiveness of targeting both juveniles and adults for reducing jellyfish abundance. The strategy to follow should depend on the biological objective. Sustaining a stable population requires adult fertile individuals, but concerns arise during summer when larger individuals pose greater risks to swimmers. If the aim is to reduce stings on bathers, the logical strategy would be the removal of juvenile jellyfish, because there will be more months with less stinging medusae. In conclusion, while removal strategies hold potential, they must balance population stability, swimmer safety, and cost considerations.

#### Change of advection rates

Advection scenarios showed the significant impact that some shoreline infrastructures could have on jellyfish populations. Breakwater construction can lead to an alteration of currents and, in turn, to an advection reduction and as a result, a local increase in jellyfish populations, as suggested by Boero (2013). A reduction in *advection rate* led to a significant increase in jellyfish abundance. These results show the importance of evaluating the effect on current patterns in the construction of breakwaters, where slight modifications of currents could lead to a dramatic increase in planktonic individuals.

#### Changes in settlement rate

Although the values of stocks and parameters for the benthic phase were estimated, this part of the model is of great value to test different hypotheses. Modification of the availability of suitable substrata to settle has shown to be of paramount importance to population stability. We showed that even a small increase or decrease in substrate availability, e.g. new breakwaters, could lead to a significant population alteration.

### Other matrix population models can be transformed into dynamic models

We showed how the modelling of a population using a matrix approach could be transformed into a dynamic model. This work outlines the steps to do it from the transition matrix values and the life cycle of the species, adding more biological reality, e.g. including the benthic phase or the graphical functions associated with death rates or the strobilation pattern, and how to optimise some parameters to obtain a good adjustment of the output to the observed data. This procedure of using matrix models to feed a more complex dynamic model could be applied to many other species already modelled using matrices.

This dynamic model allowed us to develop a model based on a transition matrix and perform an array of scenarios and analysis of sensitivity in an easier and more rapid way than in the matrix approach. The dynamic model that we built also reproduced a theoretical stable jellyfish population over 25 years, through optimising the strobilation pattern and the initial number of polyps, which allowed us to perform different sensitivity analyses and scenario comparisons.

This dynamic model provides a more accurate and comprehensive representation of the species’ life cycle, improving our understanding of its population dynamics and providing the possibility of creating an endless number of scenarios easily. We defend that dynamic models make hypothesis testing, sensitivity analysis and scenario comparison easier and more flexible than matrix models. We think that this approach has great potential to be used not only in new time-series data analysis but also in revisiting published matrix models, not only to get insights into the population ecology of the species but also to be able to run complex scenarios, e.g. related to nutrient enrichment, blooming events, harvesting, or species conservation.

## FUNDING

This work was supported by the project GVA-THINKINAZUL/2021 “A comprehensive marine observatory in the coast of Oliva-Dénia-Jávea for the conservation of biodiversity, observation of global change and promotion of the blue economy (OBSERMAR-CV).” to CB, supported by the Ministerio de Ciencia e Innovación of the Spanish Government and by Generalitat Valenciana-Conselleria de Innovación, Universidades, Ciencia y Sociedad Digital with the funding from European Union NextGenerationEU (PRTR-C17.I1).

Part of this research was conducted at the Marine Laboratory UA-Dénia https://web.ua.es/en/marlabdenia/ (Agreement Ajuntament de Dénia-Conselleria de Medio Ambiente, Agua, Infraestructuras y Territorio de la Generalitat Valenciana).

